# Seedling performance in a dioecious tree species is similar near female and male conspecific adults despite differences in colonization by arbuscular mycorrhizal fungi

**DOI:** 10.1101/2023.01.13.523901

**Authors:** Jenalle L. Eck, Camille S. Delavaux, Dara M. Wilson, Simon A. Queenborough, Liza S. Comita

**Affiliations:** Yale School of the Environment, New Haven, CT 06511, USA; Smithsonian Tropical Research Institute, Balboa, Ancón, Republic of Panama; Department of Evolution, Ecology and Organismal Biology, The Ohio State University, Columbus, OH 43210, USA; Department of Environmental Systems Science, ETH Zurich, 8092 Zurich, Switzerland

**Keywords:** plant-soil feedback, Janzen-Connell hypothesis, negative conspecific density dependence, intraspecific variation, plant-mycorrhizal mutualism, seed production

## Abstract

Plant-soil feedbacks are a key driver of species diversity and composition in plant communities worldwide; however, the factors that may cause feedbacks to vary within species are rarely examined. In dioecious species, the strength of feedbacks may differ near female plants that produce seed versus near male plants (which do not) because repeated inputs of seeds and high seedling densities near females may cause accumulation of host-specific soil microbes that influence seedling performance. To test whether conspecific seedling performance is reduced near seed-producing female trees relative to male or heterospecific trees, we conducted shadehouse and field experiments with a dioecious tropical tree species, *Virola surinamensis* (Myristicaceae), on Barro Colorado Island, Panama. The shadehouse experiment isolated the effect of soil microbial communities on seedling growth and allowed us to quantify colonization by mutualistic arbuscular mycorrhizal (AM) fungi, while the field experiment allowed us to assess seedling survival and growth in the presence of nearby conspecific adults and seedlings. In both experiments, seedling performance was similar between seedlings grown in the soil microbial communities and field environments underneath female conspecific, male conspecific, and heterospecific trees. However, contrary to expectation, seedling colonization by AM fungi was higher in male conspecific soil microbial communities than in female or heterospecific soil microbial communities at the end of the shadehouse experiment. Together, our experiments show that while differences among female and male plants in dioecious species may influence the association of conspecific seedlings with AM fungi in their soils, this variation does not necessarily translate directly to differences in seedling performance, at least over the time frame of our experiments. Studies of additional dioecious species are needed to help determine differences in soil microbial communities beneath male and female plants and to assess the role of seed input versus adult root systems in driving plant-soil feedbacks.

## Introduction

Plant-soil feedbacks are a key mechanism shaping the abundance, diversity, and composition of plant species in ecological communities worldwide (reviewed by Bever, 2003; Bonanomi et al., 2005; Ehrenfeld et al., 2005; Kulmatiski et al., 2008; Bever et al., 2010; van der Putten et al., 2013; Crawford et al., 2019), but certain aspects of their generation remain unclear. Negative plant-soil feedbacks (PSFs) are characterized by reductions in the performance of seedlings in soils conditioned by conspecific plants relative to heterospecific plants (Bever, 1994; Bever, 1997) and have been documented in a plethora of experimental studies (see meta-analyses by Kulmatiski et al., 2008; Suding et al., 2013; Crawford et al., 2019, Beals et al., 2020; Hassan et al., 2022). Negative PSFs are thought to help maintain plant species diversity by creating the patterns of seedling recruitment predicted by the Janzen-Connell hypothesis (i.e., reduced seedling recruitment nearer to adult conspecific plants or in areas of high conspecific density; Janzen, 1970; Connell, 1971; reviewed by Chesson, 2000; Hyatt et al., 2003; Carson & Schnitzer, 2008; Comita et al., 2014) and underlying the conspecific negative density-dependent dynamics that have been observed in plant species in many communities worldwide (reviewed by LaManna et al., 2017; Hülsmann et al., 2021; Song et al., 2021). Negative PSFs are generated by the accumulation of host-specific pathogenic microbes in the soil surrounding adult plants (Mills & Bever, 1998; Packer & Clay, 2003; Reinhart et al., 2005; Bell et al., 2006; Reinhart & Clay, 2009; Kotanen, 2010; Bagchi et al., 2014; Bever et al., 2015; Liang et al., 2016; Chen et al., 2019; reviewed by Song & Corlett, 2021). The presence of conspecific seeds and seedlings has also been shown to contribute to density-dependent mortality in tropical tree seedlings (Harms et al., 2000; Wright et al., 2005; Queenborough et al., 2007a; Comita et al., 2010; Metz et al., 2010; Bagchi et al., 2011; Lebrija-Trejos et al., 2014). However, because high densities of conspecific seedlings often occur near large reproductive plants, it is challenging to tease apart the influence of the adult plants versus the high density of seedlings beneath their canopy in driving pathogen accumulation and negative PSFs (but see Chanthorn et al., 2013). Experiments with dioecious plant species provide a way to separate the influence of large adult conspecifics versus the density of conspecific seeds and seedlings in driving negative PSFs.

Dioecious reproduction is exhibited by 6 % of angiosperm species globally and more than 20 % of tree species in some tropical forests (Bawa & Opler, 1975; Renner & Ricklefs, 1995; Renner, 2014), but has rarely been considered in the context of plant-soil feedbacks (Hood et al., 2004). Dioecy is characterized by separation of seed production between sexes, where female plants produce seeds while male plants do not. Sex has been shown to affect plant microbiome composition and interactions with microbes in at least some dioecious plant species (Vega-Frutis et al., 2013; Wei & Ashman, 2018; Wu et al., 2019; Guo et al., 2021; Guo et al., 2022; Li et al., 2023). Because many seeds fall beneath or close to their parent tree (Swamy et al., 2011) and seed production can vary widely among individuals even within hermaphroditic species (Hubbell, 1980), stark differences in seed and seedling input over time could result in greater pathogen accumulation in the soil surrounding female plants relative to male plants of similar size or age. This should result in decreased survival or growth of seedlings growing in the soils and environments near female plants relative to male plants, but this hypothesis has rarely been tested (but see Hood et al., 2004). Plant-soil feedback could also provide insights into one of the central paradoxes of dioecy: how do dioecious species evolve and coexist with hermaphroditic species, when only half of the individuals in their populations produce seeds (Bawa, 1980; Thomson & Barrett 1981; Käfer et al., 2017)? For dioecious species to persist in plant communities over evolutionary time, some advantages of dioecy are needed to offset the reproductive costs (Queenborough et al., 2007b, 2009). Though outbreeding and sexual dimorphism are thought to be advantages of dioecy (Smouse, 1971; Freeman et al., 1997; Zhang et al., 2023), variation in the microbiome and strength of PSFs between male and female plants is an unexplored potential benefit. If lack of seed production results in reduced conspecific seedling densities and soil pathogen accumulation near male trees, this could provide dioecious species with more enemy-free sites during seed dispersal. Variation in conspecific site suitability between males and females could lessen the overall impact of negative frequency-dependent dynamics in dioecious species (Queenborough et al., 2009), enhancing their persistence.

In addition to pathogenic microbes, co-occurring mutualists are also important in generating PSFs (Bever, 2002; Mangan et al., 2010a; Bachelot et al., 2015; Teste et al., 2017; reviewed by Revillini et al., 2016) but are underexplored in relation to dioecy. Arbuscular mycorrhizal (AM) and ectomycorrhizal (ECM) fungi can have variable effects on conspecific seedling growth and species coexistence (Bever, 2002; Castelli & Casper, 2003; Liang et al., 2015; Bennett et al., 2017; Koziol & Bever, 2018; Liang et al., 2020, Jevon et al., 2022). Positive PSFs, often caused by the accumulation of mutualistic microbes such as arbuscular mycorrhizal (AM) and ectomycorrhizal (EM) fungi in the soils near adult plants, result in increased performance of seedlings in conspecific soils relative to heterospecific soils (Smith & Reynolds, 2012; Reinhart et al., 2012; Teste et al., 2017; Segnitz et al., 2020; Duell et al., 2023). For example, positive effects due to AM fungal association may help seedlings counter conspecific distance- and density-dependent mortality due to pathogens (Liang et al., 2015; Bachelot et al., 2017; reviewed by Zahra et al., 2021), via either increases in plant growth or nutritional status (Smith & Read, 2008; reviewed by Delavaux et al., 2017) or activation of plant defensive pathways (Newsham et al., 1995; Pozo & Azcón-Aguilar, 2007). Some, but not all, dioecious and gynodioecious species exhibit sex-specific responses to mycorrhiza (Varga & Kytöviita, 2008; Eppley et al., 2009; Varga & Kytöviita, 2010a, 2010b; González et al., 2015; reviewed by Vega-Frutis et al., 2013); thus, it’s unclear whether mycorrhizal abundance, composition, and seedling colonization typically varies in the soil beneath male and female plants (Hood et al. 2004; Li et al., 2020; Lin et al., 2023) or translates to differences in seedling performance (Varga & Kytöviita 2010b; Varga et al., 2013). It’s also unclear if mycorrhizal association is or more strongly influenced by the density of conspecific seedlings or the presence of large conspecific plants. If the presence of conspecific seeds and seedlings is more important, higher mycorrhizal colonization rates would be expected in seedlings growing in female soils. In contrast, if adult presence is more important, release from competition with pathogens for root colonization sites could lead to higher mycorrhizal colonization rates in seedlings growing in male soils.

Despite evidence of the importance of PSFs in determining species composition in plant communities (reviewed by Bever, 2003; Bonanomi et al., 2005; Ehrenfeld et al., 2005; Kulmatiski et al., 2008; Bever et al., 2010; van der Putten et al., 2013; Crawford et al., 2019), it remains unclear whether the presence of large, conspecific trees or high densities of conspecific seedlings are primarily responsible for driving PSFs. To address this, we conducted shadehouse and field experiments with a dioecious tropical tree species, *Virola surinamensis* (Myristicaceae), on Barro Colorado Island, Panama. The shadehouse experiment allowed us to isolate the effect of soil microbial communities on seedling performance, while the field experiment allowed us to assess seedling performance in female and male conspecific soils in an ecologically relevant context. We asked: does seedling performance and colonization by mutualistic AM fungi vary in the soils associated with female conspecific trees versus male conspecific trees? We hypothesized that: i) seedling growth and survival would be reduced in the soils of female conspecifics, because repeated inputs of conspecific seeds and/or seedlings into female soils over time should cause greater accumulation of host-specific soil pathogens, and ii) seedling colonization by AM fungi would be higher in female soils, because the repeated input of conspecific seeds and/or seedlings over time in their soils present resources for AM fungal growth. With this study, we aim to better understand the sources of intraspecific variation in plant–soil feedbacks, allowing better predictions of how these phenomena influence abundance and composition in plant communities.

## Material and Methods

### Study site and species

Our study focused on *Virola surinamensis* (Rol. ex Rottb.) Warb. (Myristicaceae), a dioecious tropical tree species occurring on Barro Colorado Island (BCI), Republic of Panama (9°09’ N, 79°51’ W). BCI is a 15.6 km^2^ moist tropical lowland forest (Croat, 1978) receiving ∼2600 mm of rainfall per year (punctuated by a distinct dry season; Windsor, 1990). *Virola surinamensis* is native to tropical and subtropical wet lowland forests in Central America and Amazonia. It is a shade-tolerant (Howe, 1990), drought-sensitive (Fisher *et al*., 1991) canopy tree, and occurs commonly in slope and stream habitats on BCI (Harms *et al*., 2001). The minimum reproductive size for *V. surinamensis* is ∼ 30 cm diameter at 1.3 m above ground (DBH) (Riba-Hernández *et al*. 2014). Adult *V. surinamensis* are spatially aggregated but the sexes are distributed randomly relative to one another (Condit *et al*., 2000; Riba-Hernández *et al*. 2014). Flowers are fly pollinated (Jardim & Mota, 2007). On BCI, flowering occurs in the dry season (∼ Jan.) and seed production in the following wet season (∼ July) (Zimmerman *et al*., 2007). Seeds are large (∼ 2 cm) and borne singly inside a heavy, inedible capsule: ∼ 62 % of seeds are dispersed away from the maternal tree by animals (Howe & Vande Kerckhove, 1981). Herbivores predate many seeds and/or seedlings below maternal crowns, but seedlings experience higher survival if dispersed > 20 m away from the maternal tree (Howe, 1990). *Virola surinamensis* was selected from amongst the dioecious species on BCI because seed was available at the right time and germinated well enough in the shadehouse to allow adequate replication in the experiments. *Virola surinamensis* has been shown to exhibit negative PSF with some co-occurring tree species on BCI (Mangan et al. 2010b).

### Soil inoculation shadehouse experiment

To isolate the effect of soil microbes on the performance of seedlings near female and male conspecific trees, we conducted a soil microbial inoculation experiment in a shadehouse on BCI. Seeds were collected from beneath the canopy of 11 fruiting (female) *V. surinamensis* on BCI during the peak of the species’ fruiting in 2014 (between June – July). We also selected five reproductive-size male *V. surinamensis* (because flowers could not be sexed in the field, maleness was assumed because of lack of seed production during the study years, 2014 – 2016). All trees were 50 – 100 cm DBH and were located by exploring a ∼ 3.5 km^2^ area of BCI (see Fig. 1A for a map). Selected trees were located ∼ 30 m to ∼ 2 km apart. Seeds from each maternal source were surface sterilized (10 % bleach for 1 m, rinse, 70 % ethanol for 30 s, rinse), air dried, and germinated in a shadehouse in autoclaved BCI soil (collected from the forest edge near the shadehouse). One-month post-germination, a minimum of 12 healthy seedlings per maternal seed source were randomly selected for inclusion in the experiment (more seedlings were selected if available).

**Figure 1:**
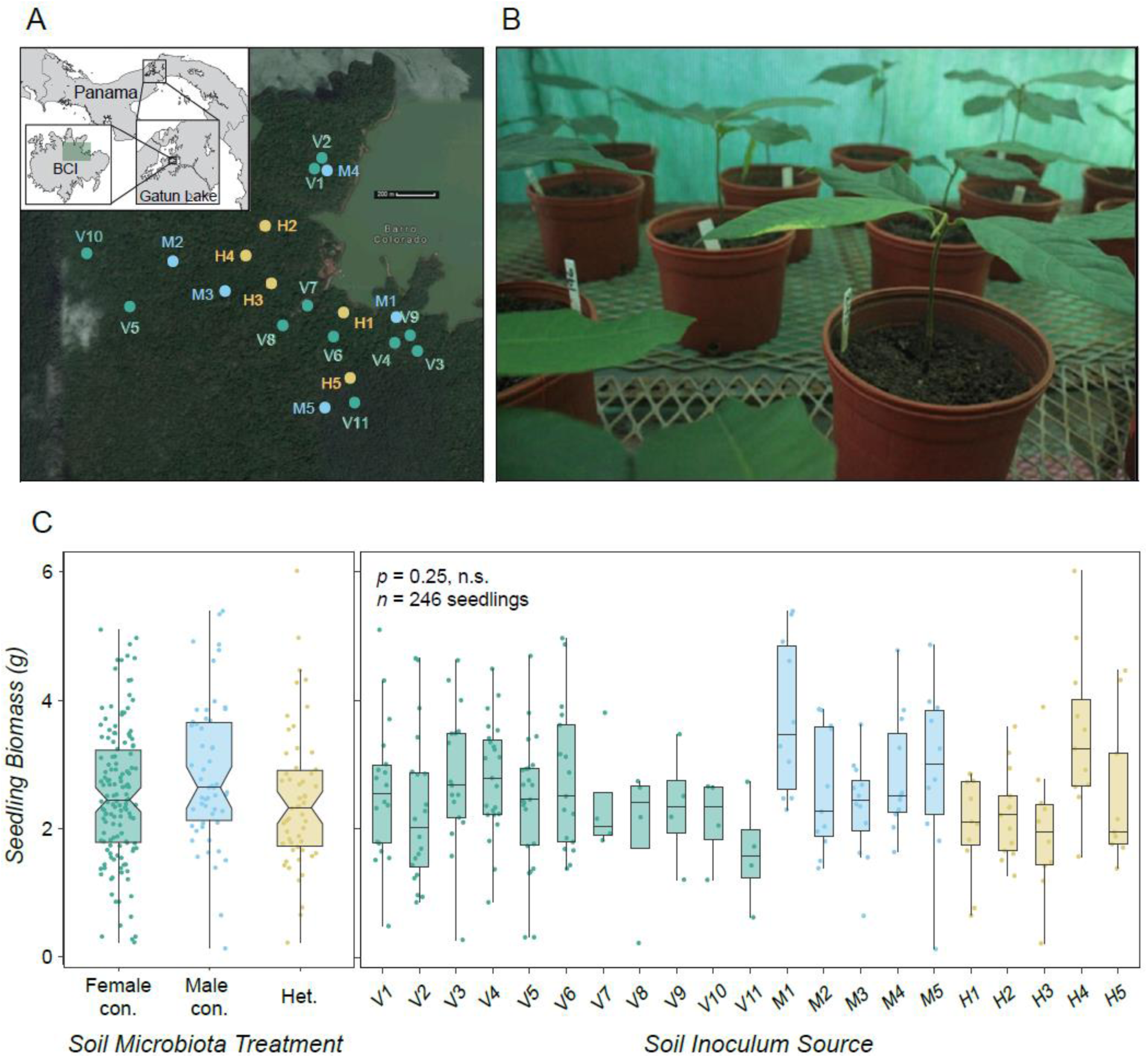
In a shadehouse experiment, seedling biomass was similar after growth in female conspecific and male conspecific soil microbial communities. *Panel A: Virola surinamensis* and heterospecific trees used as soil inoculum sources in the shadehouse experiment were located on Barro Colorado Island (BCI), Panama (the study area is shaded in green on the BCI map). Female *V. surinamensis* appear in green (V1 - V15), male *V. surinamensis* in blue (M1 - M5), and heterospecific trees in yellow (H1 – H5). Map data: Google, ©2017. Map inset modified from Baldeck *et al*., 2014. *Panel B:* In a shadehouse on BCI, we planted *V. surinamensis* seedlings in three soil microbial community inoculation treatments: female conspecific soil, male conspecific soil, or heterospecific soil. *Panel C*: At the end of the 8-mo shadehouse experiment, the biomass of *V. surinamensis* seedlings grown in the soil microbial community associated with female conspecific trees was similar to seedlings grown in the soil microbial community associated with male conspecific trees or heterospecific trees (Table S1). In *C,* box belts show the median values, box notches represent a 95 % confidence interval for comparing medians, box hinges correspond to the first and third quartiles, and box whiskers extend to the largest and smallest value no further than 1.5 × the interquartile range from the hinges.

We collected samples of the soil microbial community beneath the canopy of each of the source trees to use as inocula in the experiment. To create one soil microbial inoculum per tree, we collected 2 L of soil at a depth of 10 cm from three randomly selected points within 3 m of each trunk (to roughly correspond to fine roots, where soil microbes are likely to be most active), then coarsely sieved and homogenized the soil from these three points. To act as a control, we also created soil microbial inocula from beneath the canopy of five reproductive size heterospecific trees within the study area (Fig. 1A). Heterospecific trees (abbreviated as H1 – H5) were chosen to represent a range of common species and families (H1: *Spondias mombin* L. (Anacardiaceae); H2: *Ormosia macrocalyx* Ducke (Fabaceae); H3: *Anacardium excelsum* (Bertero & Balb. ex Kunth) Skeels (Anacardiaceae); H4: *Platypodium elegans* Vogel (Fabaceae); H5: *Protium tenuifolium* (Engl.) Engl. (Burseraceae)). Each seedling was assigned to the soil inoculum of only one of the 21 adult trees in the study (i.e., the 11 female conspecific trees, five male conspecific trees, or five heterospecific trees). Thus, seedlings from each maternal seed source were assigned at random to each of three soil inoculum treatments: female conspecific soil inoculum (*n* = 144 seedlings), male conspecific soil inoculum (*n* = 60 seedlings), and heterospecific soil inoculum (*n* = 57 seedlings). More seedlings were planted in female soil inoculum because the experiment was designed to test concurrently for the effect of relatedness on seedling performance in maternal soils (see Eck et al., 2019). All soil inocula were collected in September 2014 and used in the experiment within ∼ 3 days.

In a shadehouse on BCI, 261 experimental seedlings were transplanted into individual 2-L pots in September 2014 (at ∼ 1–2 mo. of age). Each pot contained 20 % by volume of soil microbial inoculum from the seedling’s assigned tree and 80 % by volume of a common soil medium (a steam-sterilized 1:1 mixture of BCI field soil:potting sand). To inoculate the seedlings, each pot was filled 2/3 of the way with common medium and a circular depression created in the middle, into which the soil inoculum was placed. Seedlings were transplanted directly into the inoculum and then a top layer of common medium was placed to cover the inoculum and roots. This ensured that the seedling roots encountered the soil inoculum early in the experiment (as experimental time was limited, relative to tree life history spans) while having access to nutrients in the common medium for further growth. Relatively small volumes of field soil inoculum were used to minimize the potential impact of differences in soil nutrients among inocula on seedling performance. Seedling pots were placed across four shadehouse benches in a balanced, randomized design (see Fig. 1B for a photograph of the shadehouse experiment). Shadehouse benches were covered with two layers of 80 % shade cloth (to mimic shady understory conditions) and were shielded from rainfall with a roof of clear plastic lining. Stem height, the number of leaves, and the length and width of each leaf were measured for each seedling immediately after transplant and were measured again periodically throughout the experiment (along with survival). Initial oven-dried biomass was estimated for each experimental seedling based on stem height during the first census using an allometric linear regression model (F_(1,42)_ = 338.1, *p* < 0.001, R^2^ = 0.887; Eck et al., 2019). This model was built based on measurements of height and dry biomass of a randomly harvested sample of potential experimental seedlings at the beginning of the experiment. Seedlings remained in their experimental treatments for ∼ 8 mo. and were amply watered three times per week.

In May 2015, the 250 surviving seedlings were harvested, and total oven-dried biomass (including aboveground and belowground portions), stem height, the number of leaves, and total leaf area were measured for each seedling. To quantify colonization by AM fungi in the surviving seedlings, we used the magnified root intersect method (McGonigle *et al*., 1990) described in Eck *et al*. 2019 (Supplementary Methods). To more fully assess the potential factors influencing seedling performance in the shadehouse and test whether nutrient availability during the experiment was linked to seedling performance, we also quantified soil nutrient conditions in each soil inocula at the end of the experiment using the method described in Eck *et al*. 2019 (Supplementary Methods).

### Field experiment

To test whether seedling performance was reduced in the soils beneath female conspecific trees relative to male conspecific and heterospecific trees in an ecologically relevant setting, we conducted a field experiment in the BCI forest. We collected seeds from beneath the canopy of six female *V. surinamensis* during the June–July 2015 fruiting season. Because of seed availability and germination rates, only two of the female trees from the shadehouse experiment were also used in the field experiment (V5 and V11). Additional female trees were located by exploring the same ∼ 3.5 km^2^ area of BCI as in the shadehouse study (see Fig. 3A for a map and Fig. 3B for a photograph of a focal *V. surinamensis*). All five male *V. surinamensis* and four of the heterospecific trees studied in the shadehouse experiment were included in the field experiment (H2, *Ormosia macrocalyx,* was replaced with a new heterospecific tree, H6: *Sterculia apetala* (Jacq.) H. Karst (Malvaceae)). Seeds from each maternal source were surface-sterilized, air dried, and germinated in autoclaved BCI forest soil in a shadehouse under two layers of 80 % shadecloth.

**Figure 2.**
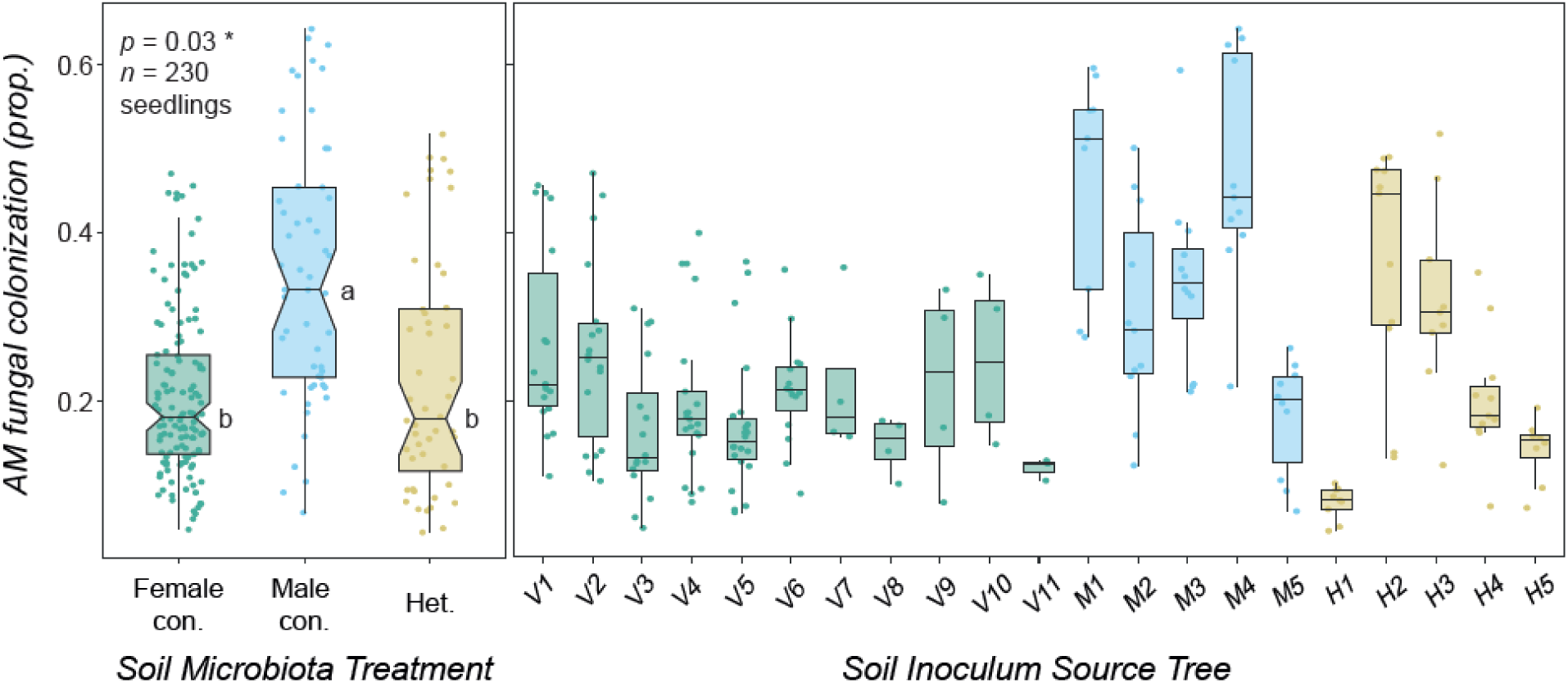
Seedling colonization by AM fungi was higher after growth in the soil microbial communities associated with male conspecific trees in a shadehouse experiment. At the end of the 8-mo shadehouse experiment on Barro Colorado Island (Panama), colonization by AM fungi was higher in *Virola surinamensis* seedlings grown in the soil microbial communities associated with male conspecific trees relative to seedlings grown in soil microbial communities associated with female conspecific trees or heterospecific trees (Table S2). In four of the five male soil microbial communities tested, the median predicted values for AM fungal colonization were higher than in any of the 11 female soil microbial communities tested. Box belts show the median predicted values, box hinges correspond to the first and third quartiles, while box whiskers extend to the largest and smallest value no further than 1.5 × the interquartile range from the hinges (predicted values are plotted to show patterns after accounting for covariates). Box notches represent a 95 % confidence interval for comparing predicted medians.

**Figure 3.**
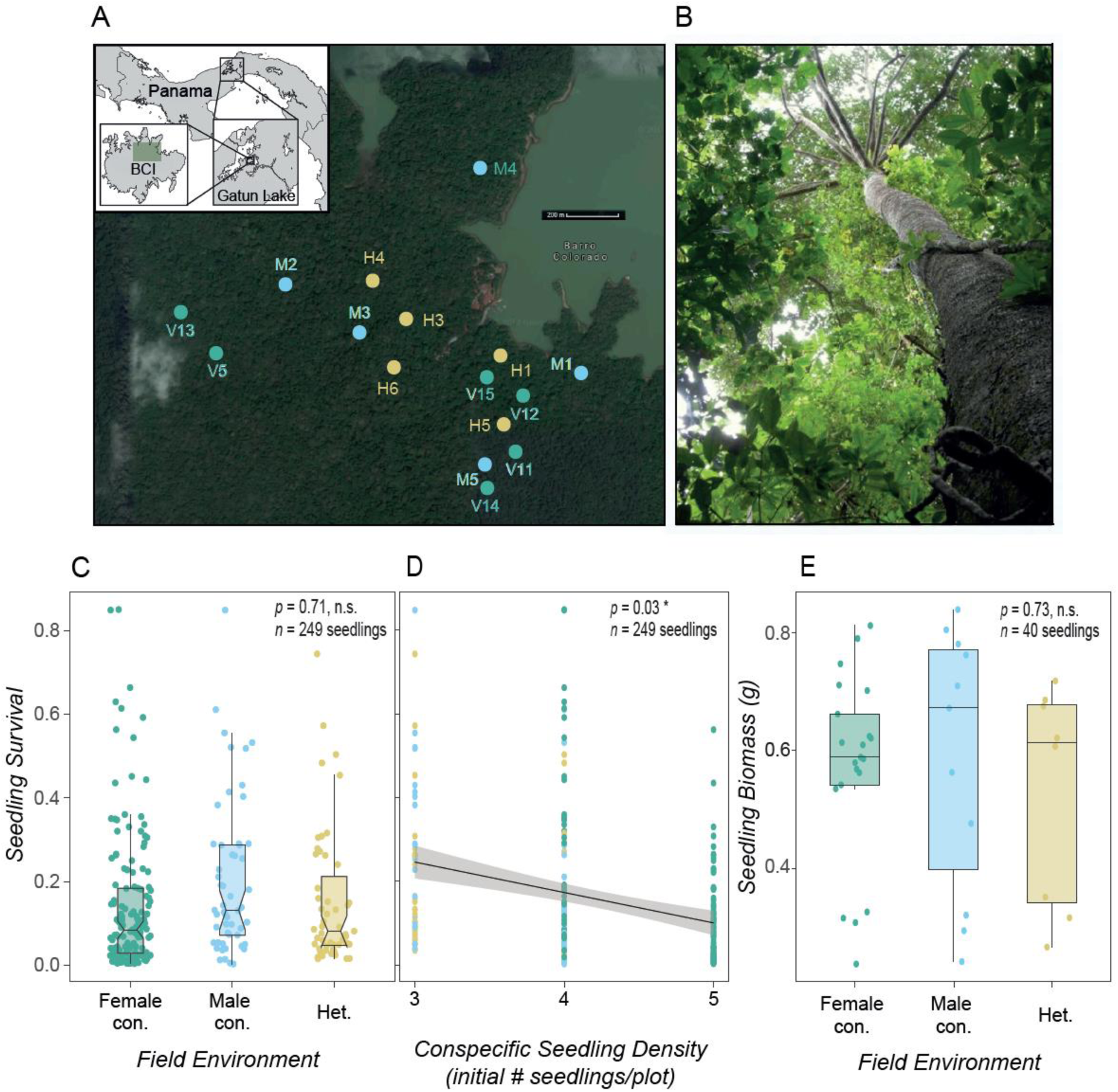
In a field experiment, seedling performance was similar in female conspecific and male conspecific environments but was influenced by conspecific seedling density. *Panel A: Virola surinamensis* and heterospecific trees used as soil inoculum sources in the field experiment are located on Barro Colorado Island (BCI), Panama (the study area is shaded in green on the BCI map). Female *V. surinamensis* are marked in green (V5, V11 - V15), male *V. surinamensis* in blue (M1 - M5), and heterospecific trees in yellow (H1, H3 – H6). Map data: Google, ©2017. Map inset modified from Baldeck *et al*., 2014. *Panel B:* In a field experiment on BCI, we planted *V. surinamensis* seedlings in three field treatments: female conspecific environments, male conspecific environments, or heterospecific environments. *Panel C*: At the end of the 7-mo field experiment, the survival of *V. surinamensis* seedlings was similar among female conspecific, male conspecific, and heterospecific field environments (Table S3). *Panel D*: Seedling survival at the end of the field experiment decreased as the number of experimental conspecific seedlings initially placed in the field plots increased (Table S3). *Panel E:* The biomass of the *V. surinamensis* seedlings that survived until the end of the experiment was similar among seedlings growing in female conspecific, male conspecific, and heterospecific field environments (Table S4). In *C* & *E*, predicted values are plotted as points, box belts show the median predicted values, box notches represent a 95 % confidence interval for comparing predicted medians, box hinges correspond to the first and third quartiles, while box whiskers extend to the largest and smallest value no further than 1.5 × the interquartile range from the hinges. In *D*, the shaded area is a 95 % confidence interval band surrounding the best fit regression line.

One-month post-germination, a minimum of 18 healthy seedlings per maternal seed source were randomly selected for inclusion in the field experiment. Within each seed source, seedlings were randomly assigned to one of three experimental field soil treatment groups: beneath a male conspecific adult, beneath a female conspecific adult, or beneath a heterospecific adult. Seedlings were randomly assigned to be transplanted beneath one adult tree within their treatment group, and into one of 3–6 seedling plots (1 m^2^) beneath the canopy of their assigned tree. Plot locations were randomized with respect to direction and distance from the base of the focal tree (1–4 m). Plots contained 3–5 experimental seedlings, each of which were randomly assigned to a 25-cm^2^ position within the plot. Because the field experiment was designed to concurrently test for the effect of relatedness on seedling performance near maternal trees (Eck et al., 2019; Eck et al., in prep.), female trees had, on average, more plots per tree and more seedlings per plot than male trees.

The 249 experimental seedlings were transplanted at ∼ 1 mo of age into their field treatment plots. Seedlings roots were rinsed gently to remove excess potting soil before transplant. Immediately after transplant (August 2015), stem height, the number of leaves, and the length and width of each leaf were measured for each seedling. Each seedling was also stem-tagged with a unique identification number. All seedlings were censused every 1–2 mo, during which time survival was recorded, and the stem height and number of leaves were measured for all surviving seedlings. We also recorded instances of seedling stem breakage and uprooting (likely caused by mammalian or avian herbivores). Seedlings were not watered during the field experiment and received only ambient rainfall. Seedlings that disappeared (with or without their tags being found) were recorded as dead during the census of their disappearance. After ∼ 7 mo in the field experimental treatments (late March 2016), the 40 surviving seedlings were harvested and their total oven-dried biomass (including aboveground and belowground portions), stem height, and total leaf area were measured.

### Statistical analyses

#### Seedling performance in the shadehouse experiment

To test whether seedling performance was reduced in female conspecific soil microbial communities relative to male conspecific and heterospecific soil microbial communities in the shadehouse experiment, we built a linear mixed-effects model (LMM). We focused on growth (i.e., biomass) of seedlings that survived until the end of the experiment because survival rates in the shadehouse were high (95.8 %; Supplementary Methods). We modelled seedling oven-dried biomass at harvest in all conspecific and heterospecific soils. Fixed effects in this model included soil microbial treatment (female conspecific, male conspecific, or heterospecific), estimated initial seedling biomass, soil nutrient conditions, and the size (DBH) of the soil inoculum source tree. Maternal seed source, soil inoculum source, and shadehouse bench were included as crossed random effects. Seedlings without initial or final biomass data were excluded from all models. All statistical analyses in our study were conducted in the R statistical environment (R Core Team 2022). Linear mixed-effects models in our study were constructed using the lme4 package (Bates et al., 2015). P values for LMM predictors were obtained using the lmerTest package (Kuznetsova et al., 2016).

#### Seedling colonization by AM fungi in the shadehouse experiment

To test whether seedling colonization by AM fungi in the shadehouse experiment was higher in female conspecific soil microbial communities relative to male conspecific and heterospecific soil microbial communities, we built a generalized linear mixed-effects model (GLMM). The proportion of root intersects colonized by any AM fungal structure was modelled in each seedling that survived until the end of the shadehouse experiment using the glmmTMB package (Brooks et al., 2017). Though we focus here on analyses utilizing data on all visible AM fungal structures, we also examined the proportion of root intersects colonized by i) arbuscules and ii) vesicles (Supplementary Information). Fixed effects in the model utilizing all AM fungal structures included soil microbial treatment, estimated initial seedling biomass, soil nutrient conditions, and the size (DBH) of the soil inoculum source tree. To account for potential variation between the two researchers that quantified AM fungal colonization in the seedlings, we also included observer as a fixed effect in the model. Random effects in the model included maternal seed source, soil inoculum source, and shadehouse bench. Model family was set to beta-binomial (to counter overdispersion) and weights were set to the number of root intersects quantified in each seedling. P values for GLMM predictors were obtained using the car package (Fox & Weisberg, 2019).

#### Seedling performance in the field experiment

To test whether seedling performance in the field experiment was reduced in female conspecific environments relative to male conspecific and heterospecific environments, we built a series of mixed-effects models analyzing seedling survival and growth. First, the survival of seedlings in all conspecific and heterospecific environments was modelled at the end of the field experiment as a binary response variable using a GLMM with binomial errors. Fixed effects in the model included field environment (female conspecific, male conspecific, or heterospecific), estimated initial seedling biomass, and the size (DBH) of the nearby adult tree. We also included whether a seedling experienced stem breakage and/or uprooting (at any point during the field experiment) as a binary fixed effect in the model. Furthermore, we also included the number of experimental seedlings in the field plot as a fixed effect in this model. Maternal seed source and the unique identifier of the adult tree each seedling was planted near were included as random effects.

Next, we tested whether the biomass of the seedlings that survived until the end of the field experiment varied among female conspecific, male conspecific, and heterospecific field environments. We modelled oven-dried harvest biomass with a separate LMM containing the same set of fixed and random effects specified in the prior survival model. We also tested for differences in seedling growth during the experiment with a LMM using the relative growth rate (RGR) of each living seedling during each census interval as the response variable (utilizing 350 total observations of 101 individual seedlings). Relative growth rates were calculated based on field measurements of seedling stem height and leaf number (Equation 1). Field measurements were converted to estimated seedling biomass (*S*) during each of census (*t*) using the aforementioned allometric model.

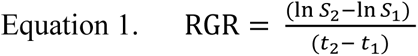

The RGR model included the set of fixed effects specified for the prior survival model. Random effects in the model included maternal seed source, the unique identifier of the adult tree each seedling was planted near, the unique identifier of the seedling (to account for repeated measurements of seedlings over time), and census interval. In both models, random effects that explained zero variance were removed from the final models.

## Results

### Seedling performance in the shadehouse experiment

At the end of an 8-mo shadehouse experiment, we found that *V. surinamensis* seedlings grown in a soil microbial community associated with a female conspecific tree had similar biomass as seedlings grown in a soil microbial community associated with a male conspecific tree or a heterospecific tree (Fig. 1C & Table S1; *F* = 1.52, *p* = 0.25; *n* = 246 seedlings). Seedlings grew substantially during the experiment: the average biomass of seedlings at the end of the experiment was 2.58 ± 1.08 g versus an estimated 0.40 ± 0.14 g at the beginning of the experiment. We found no relationship between seedling biomass and the size of the conspecific soil inoculum source tree (Fig. 1D & Table S1; *F* = 1.64, *p* = 0.22; *n* = 246 seedlings). or nutrient conditions in the soil inocula at the end of the experiment (Table S1; *F* = 1.93, *p* = 0.18; *n* = 246 seedlings).

### Seedling colonization by AM fungi in the shadehouse experiment

Most of the seedlings (82.5 %) were colonized by AM fungi after 8-mo. of growth in the shadehouse experiment. We found that seedlings grown in a soil microbial community associated with a male conspecific tree had higher proportions of colonization by any AM fungal structure in their roots than seedlings grown in a soil microbial community associated with a female conspecific tree or a heterospecific tree (Fig. 2 & Table S2; *p* = 0.03; *n* = 230 seedlings). This result was consistent across soil inoculum sources: in four out of the five male *V. surinamensis* trees in our study, the predicted mean proportions of seedling colonization by AM fungi were higher than the means predicted for any of the 11 female *V. surinamensis* trees. When examining specific AM fungal structures, seedlings in male conspecific soils also had higher proportions of colonization by arbuscules relative to seedlings in female soils (Table S2; *p* = 0.02, *n* = 230 seedlings), while proportions of seedling colonization by vesicles were similar in female and male conspecific soils (Table S2; *p* = 0.29, *n* = 230 seedlings). We found that seedling colonization by AM fungi was not related to the size of the conspecific tree providing the soil inocula (Table S2; *p* = 0.25; *n* = 230 seedlings) or soil nutrient conditions in pots at the end of the experiment (Table S2; *p* = 0.47; *n* = 230 seedlings).

### Seedling performance in the field experiment

At the end of the field experiment (Figs. 3A & 3B), we found that seedling survival was similar among seedlings planted in female conspecific environments, male conspecific environments, and heterospecific environments in the field (Fig. 3C & Table S3; *p* = 0.71; *n* = 249 seedlings). Seedling mortality was high with only 16.1 % of the experimental seedlings alive after 7 mo. We found that seedlings in field plots that contained higher densities of experimental seedlings at the beginning of the experiment had lower survival (Fig. 3D & Table S3; *p* = 0.03; *n* = 249 seedlings). The size of the focal tree the seedling was planted near did not affect seedling survival (Table S3; *p* = 0.67, *n* = 239 seedlings).

As with survival, the harvest biomass of seedlings that survived until the end of the field experiment was similar among female conspecific, male conspecific, and heterospecific field environments (Fig. 3E & Table S4; *F* = 0.33; *p* = 0.73; *n* = 40 seedlings). Seedling biomass at the end of the field experiment was not related to the initial density of experimental seedlings in the field plots (Table S4; *F* = 1.62; *p* = 0.21; *n* = 40 seedlings) or to the size of the focal tree the seedling was planted near (Table S4; *F* = 1.19; *p* = 0.29; *n* = 37 seedlings). Similar results were found when analyzing seedling relative growth rates during the experiment with a repeated measures model (Table S5).

## Discussion

Though recent studies have focused on quantifying patterns of, and variation in, plant-soil feedback among co-occurring species (reviewed by Smith-Ramesh & Reynolds, 2017; Crawford et al., 2019; Bennett & Klironomos, 2019, Beals et al., 2020), less is known about the causes and consequences of intraspecific variation in PSFs (Felker-Quinn et al., 2011; Hovatter et al., 2013; Bukowski & Petermann, 2014; Liu et al., 2015; Wagg et al., 2015; Allen et al., 2018; Florianová & Münzbergová, 2018; Eck et al., 2019; Crawford & Hawkes, 2020; Rallo et al., 2023). Differences in seed production among adult plants of dioecious species offers the opportunity to disentangle the effects of adult root systems from the input of conspecific seeds and seedlings in driving PSFs. In complementary shadehouse and field experiments in Panama, we tested whether seedling performance and colonization by AM fungi differed in the soil microbial communities and field environments associated with female conspecific and male conspecific adult trees in a dioecious tropical tree species. We did not find evidence in support of the hypothesis that seedling performance is reduced in female soils relative to male soils due to higher accumulation of host-specific pathogenic microbes. Seedling survival and growth in the field and seedling growth in the shadehouse were similar among seedlings grown in female conspecific, male conspecific, and heterospecific soils. We did find that seedling colonization by AM fungi differed in male versus female soils. However, contrary to our expectation, seedling colonization by AM fungi was more frequent in the roots of seedlings grown in male conspecific soils than in the roots of seedlings grown in female conspecific or heterospecific soils. Together, our experiments show that while differences among female and male plants in dioecious species may influence the association of conspecific seedlings with AM fungi in their soils (suggesting that differences in soil microbial composition between sexes may occur in this species), this variation does not necessarily translate directly to differences in seedling performance, at least over the time frame of our experiments. Studies of additional dioecious species are needed to help determine differences in soil microbial communities beneath male and female plants and to assess the role of seed input versus adult root systems in driving plant-soil feedbacks.

Mutualistic AM fungi are thought to be important in influencing plant-soil feedbacks and seedling performance (Bever, 2002; Mangan et al., 2010a; Bachelot et al., 2015; Liang et al., 2015; Bachelot et al., 2017; Bennett et al., 2017; Jevon et al., 2022). Studies with other dioecious species show that individuals of different sexes can form different associations with AM fungi (Varga & Kytöviita, 2008, 2010a, 2010b; Eppley et al., 2009; Vega-Frutis et al., 2013), but it’s unclear what consequences this variation might have in natural plant populations. Increased AM fungal colonization in the soils beneath male trees in our study suggests that adult sex does affect interactions with at least some microbial groups in *V. surinamensis*. We hypothesized that the repeated input of conspecific seeds and/or seedlings near female trees would result in more opportunities for the growth of AM fungi, and subsequently, higher colonization rates by AM fungi in seedlings grown in female conspecific soils relative to male soils. Because colonization by AM fungi was higher in seedlings grown in male conspecific soils relative to female conspecific soils in our study, this suggests that another mechanism may be affecting AM fungal association. One potential mechanism is the release of AM fungal growth (or seedling association with AM fungi) in male soils from pathogen suppression or soil nutrient constraints in female soils. Hood et al. (2004) found differences in AM fungal colonization depending on whether seedlings were grown in conspecific versus distant soils, but not between female versus male conspecific soils, suggesting that these dynamics could vary among plant species.

In our study, differences in AM fungal colonization did not translate to differences in seedling performance in the soil microbial communities associated with male conspecific versus female conspecific trees. Though negative impacts of the soil microbial communities near adults on conspecific seedlings appear to be more common than positive impacts in AM fungal-associated trees (Mangan et al., 2010b; Liu et al., 2012; McCarthy-Neumann & Ibáñez, 2013; Bennett et al., 2017; Eck et al., 2019), AM fungal benefits to seedlings are strong enough to balance the negative impact of host-specific pathogens in at least some tree species (Mangan et al., 2010a; Liang et al., 2015). Colonization rates by AM fungi are not always linked to plant growth benefits (Graham et al., 1991; Gange & Ayres, 1999), e.g., if the colonizing AM fungal species primarily provide physical pathogen protection, rather than nutritional benefits. However, because the incidence of arbuscules (but not vesicles) differed between seedlings in male and female soils in our study, this suggests that AM fungi could function more efficiently for nutrient transfer in male soils of this species, while lipid storage in seedling roots may be similar in female and male soils (Johnson, 2010). In addition, because growth effects due to soil microbes are the net result of pathogens, mutualists, and all other drivers, similarity in conspecific seedling biomass is not necessarily indicative of the relative impact of AM fungi in male versus female soils. AM fungal-associating tree species generally show stronger negative conspecific density-dependence than EM fungal-associating tree species (Jiang et al., 2020; Delavaux et al., *in prep.*), indicating that the impact of AM fungi are relatively weak compared to pathogens in AM fungal-associated plant communities. Thus, if greater pathogen impacts occur for seedlings in female soils, mycorrhizal benefits would also need to be greater in female soils than in male soils to result in observations of similar seedling biomasses between the two. Studies that quantify pathogen and mutualist community composition, patterns of seedling infection, and mycorrhizal function alongside mycorrhizal association in male and female soils will help disentangle these effects.

The advantages and disadvantages of the dioecious reproductive system and how dioecious species are maintained in plant communities presents a paradox in ecology (Queenborough et al., 2007b, 2009; Bruijning et al., 2017; Bruns et al., 2018). Though variation in plant-pathogen interactions among sexes has been demonstrated in several dioecious species (Alexander, 1989; Åhman, 1997; Shykoff et al., 1997; Hood et al., 2004; Moritz et al., 2016) and could help drive the evolution of dioecy (Bruns et al., 2018), the relationships between negative PSFs and dioecy are just beginning to be considered (but see Hood et al., 2004). Our study adds to this discussion by suggesting that male environments may provide an additional advantage over female environments: higher colonization rates by mutualist AM fungi. Higher colonization by AM fungi (or other mutualist advantages) in some conspecific sites could cause reductions in negative frequency-dependence in dioecious species, affecting their ability to coexistence in plant communities (Montesinos et al., 2007). Though evolutionary expectations predict reduced mycorrhizal interactions in males of tropical tree species (Vega-Frutis et al., 2015), colonization rates in male soils may be the result of indirect effects of male environments (e.g., lack of seed input) rather than direct interactions between male adult trees and AM fungi. Because we did not find differences in seedling growth or survival near male and female trees, our study also suggests overall tolerance of seedlings to female seed production (and any associated environmental changes), which could help seedlings compensate for any potentially negative impacts of aggregated seed production (e.g., higher enemy densities). Similarity in seedling performance in male conspecific, female conspecific, and heterospecific environments could also help explain the aggregated patterns of local distribution that have been observed in *V. surinamensis* (Condit et al., 2000; Riba-Hernández et al., 2014).

Though negative plant-soil feedbacks have been documented in our study species (Mangan et al., 2010b), we did not find differences between *V. surinamensis* seedling performance in conspecific soils versus heterospecific soils in our study. However, because plant-soil feedbacks arise from feedback between plants and the biotic or abiotic conditions of the soil, while pairwise negative plant-soil feedbacks between co-occurring species are what is needed to assess the potential for species coexistence (rather than comparisons of the performance of a single species in conspecific versus heterospecific soils; Bever, 1997), our result does not necessarily contradict prior studies demonstrating pairwise negative plant-soil feedbacks in *V. surinamensis* (Mangan et al., 2010b). Thus, host-specific soil-borne pathogens could still be a key factor determining seedling performance in our study species (Eck et al., 2019). Because we did not quantify enemy densities in our study, we cannot rule out the possibility that seed production could cause short- or long-term changes to the density or composition of soil pathogens or other natural enemy communities near established plants. Effects of seed production on conspecific seedling performance via natural enemies (or mutualists) could still occur in other species, emerge over longer time periods or during later developmental stages, or could be critically determined during or immediately after seed germination. Though it was not the focus of this study, we found that increases in the number of experimental conspecific seedlings in our field plots negatively affected seedling survival, in line with other studies showing the importance of conspecific seedling neighbors in determining seedling performance (Harms et al., 2000; Queenborough et al., 2007a; Comita et al., 2010; Metz et al., 2010; Lebrija-Trejos et al., 2014). In at least one subtropical tree community, a positive relationship was found between the density of conspecific adults and soil pathogens that led to higher conspecific seedling mortality (Liang et al., 2016); thus, the density of conspecific adults may have a larger effect on enemy densities than the presence of just the nearest conspecific adult.

Plant genotype and the level of genetic relatedness between plants may also influence seedling performance in conspecific soils (Smith et al., 2012; Bukowski & Petermann, 2014; Browne & Karubian, 2016; Eck et al., 2019; Duell et al., 2023). In our study species, shadehouse seedlings that were grown in the soil microbial community associated with their maternal tree had lower biomass than seedlings that were grown in the soil microbial community of an unrelated conspecific female adult (Eck et al., 2019). This finding demonstrates that the shadehouse experimental approach utilized in our study can detect differences in conspecific soil effects. Together, these shadehouse results suggest that genetic relatedness to the seedling, rather than adult seed production, determines the effect of conspecific soil microbial communities on seedling performance in this species. Lack of differences in seedling growth between male and female conspecific soil microbial inoculum sources in our study could be because i) we did not account for relatedness of the soil donor tree to the focal seedling and ii) the soil donor trees likely had varying levels of relatedness to the experimental seedlings.

Predicting how and when plant-soil feedbacks occur remains a central challenge in ecology (reviewed by Bever, 2003; Kulmatiski et al., 2008; Bever et al., 2010; van der Putten et al., 2013; Crawford et al., 2019). Variation in biotic interactions among individuals within populations is often overlooked in community ecology but is an important component affecting ecological dynamics (reviewed by Freckleton & Lewis, 2006; Bolnick et al., 2011). The demographic patterns predicted by Janzen (1970) and Connell (1971) have been shown to occur in a variety of plant communities worldwide (reviewed by Carson & Schnitzer 2008; Terborgh, 2012; Comita et al., 2014), and theoretical studies demonstrate that they can enhance local species richness (Adler & Muller-Landau, 2005; but see Stump & Comita, 2018) and structure the relative abundance of species in plant communities (Mangan et al., 2010b). Recently, studies expanding on the Janzen–Connell hypothesis to consider intraspecific effects within populations have found that within-species variation may also contribute to the maintenance of species diversity and the evolution of plant traits, such as seed dispersal (Schupp et al., 1992; Liu et al., 2015; Browne & Karubian, 2016; Eck et al., 2019). Variation in PSFs within species could affect plant species diversity by enabling species to differ in the distributions of their responses to the environment (Clark, 2010) or by modulating the number of enemy-free sites during seed dispersal (Eck et al., 2019). Our study does not provide evidence of variation in PSFs within species, but instead shows variation in biotic interactions between conspecific plants and AM fungal mutualists. Understanding the factors that influence conspecific seedling survival near adult plants will help us better understand how natural enemies and mutualists structure plant diversity and abundance.

