## Supplementary Information for "Seedling performance in a dioecious tree species is similar near female and male conspecific adults despite differences in colonization by arbuscular mycorrhizal fungi"

<u>The following Supporting Information is available for this article:</u>

**Supplementary Methods.** Quantification and analysis of colonization by AM fungi in the experimental seedlings.

**Supplementary Methods.** Quantification and analysis of nutrients in the soil inocula in the shadehouse experiment

**Figure S1.** A micrograph of AM fungal structures quantified in the roots of experimental seedlings.

**Figure S2**. Loading plot of soil nutrient variable contributions and dimensionality in principal component analysis.

**Table S1**. Effect of soil microbial treatment on seedling biomass in the shadehouse experiment.

**Table S2.** Effect of soil microbial treatment on seedling colonization by AM fungi in the shadehouse experiment.

**Table S3.** Effect of female conspecific, male conspecific, and heterospecific field environments on seedling survival in the field experiment.

**Table S4.** Effect of female conspecific, male conspecific, and heterospecific field environments on seedling biomass in the field experiment.

**Table S5:** Effect of female conspecific, male conspecific, and heterospecific field environments on seedling relative growth rates in the field experiment.

**Table S6.** Summary of principal component analysis of nutrients in the soil inocula at the end of the shadehouse experiment.

**List of references cited only in the Supplementary Information.**

**Dataset S1.** Shadehouse experimental data (separate file).

**Dataset S2.** Field experimental data (separate file).

**Dataset S3**. R code for shadehouse analyses (separate file).

**Dataset S4.** R code for field analyses (separate file).

### Supplementary Methods

*Quantification and analysis of colonization by AM fungi in the shadehouse seedlings*

We quantified colonization by AM fungi in the roots of the seedlings that survived until the end of the shadehouse experiment using the method described in Eck *et al*. 2019. We obtained AM fungal colonization data for 230 of the surviving experimental seedlings. A micrograph showing the structures quantified can be viewed in Fig. S1. We calculated AM fungal colonization for each seeding as the proportion of root intersects quantified in the subsample that contained any visible AM fungal structure. As a more conservative estimate of AM colonization and to better explore potential AM fungal function, we also calculated the proportion of root intersects quantified that contained visible arbuscules only and visible vesicles only.

In addition to modelling AM fungal colonization rates among seedlings based on all AM fungal structures (as described in the statistical methods section of the main text), we also built models testing for differences in AM fungal colonization rates among seedlings based on the occurrence of i) arbuscules only and ii) vesicles only (Table S2) using the glmmTMB package (Brooks et al., 2017). Fixed effects in these models included soil microbial treatment, estimated initial seedling biomass, soil nutrient conditions, and the size (DBH) of the soil inoculum source tree. To account for potential variation between the two researchers that quantified AM fungal colonization in the seedlings, we also included observer as a fixed effect in the model. Random effects in the model included maternal seed source, soil inoculum source, and shadehouse bench. Model family was set to beta-binomial (to counter overdispersion) and weights were set to the number of root intersects quantified in each seedling. P values for GLMM predictors were obtained using the car package (Fox & Weisberg, 2019).

*Quantification and analysis of nutrients in the soil inocula in the shadehouse experiment*

Our shadehouse experiment was designed to minimize soil nutrient differences among soil microbial inoculum sources (i.e., by using only 20% by volume of field soil inoculum and 80% by volume of a common growing medium in each pot). However, because soil variables have previously been shown to be a key driver of tree species distributions and seedling growth in central Panama (Condit et al., 2013; Zalamea, 2016), we also assessed the effect of soil nutrients on seedling biomass. To test whether soil nutrient availability during the shadehouse experiment affected seedling performance, we quantified soil nutrient conditions in each of the 21 soil inocula at the end of the shadehouse experiment using the method described in Eck et al., 2019. Eight variables were quantified for each inoculum: soil pH in water, soil pH in CaCl2, total soil phosphorous (total P mg/soil kg), plant-available phosphorous in soil (Bray 1-P mg/soil kg), soil organic matter (% loss on ignition), soil carbon (% total C), soil nitrogen (% total N), and soil C : N ratio.

To reduce the dimensionality of the soil nutrient variables for explanatory use in analyses, we conducted principal component analysis (PCA) using the standardized variables. During PCA, all eight soil nutrient variables loaded on the first principal component (PC1), which explained 47 % of the variance; the second principal component (PC2) loaded six variables and explained an additional 22 % of variance (Fig. S2 and Table S6). We included soil nutrient conditions in the relevant models of seedling performance in the shadehouse experiment using the first principal component (PC1) of the PCA as a fixed explanatory variable. In addition, because nitrogen measurements might not be meaningful after soil storage, we performed an identical PCA and analyses as described above, but after excluding our nitrogen-related variables (% total N and C : N ratio). Model results were similar when nitrogen-related variables were excluded versus included, so we focus on the results of the model including nitrogen, utilizing all available soil information.

**Supplementary Figures**

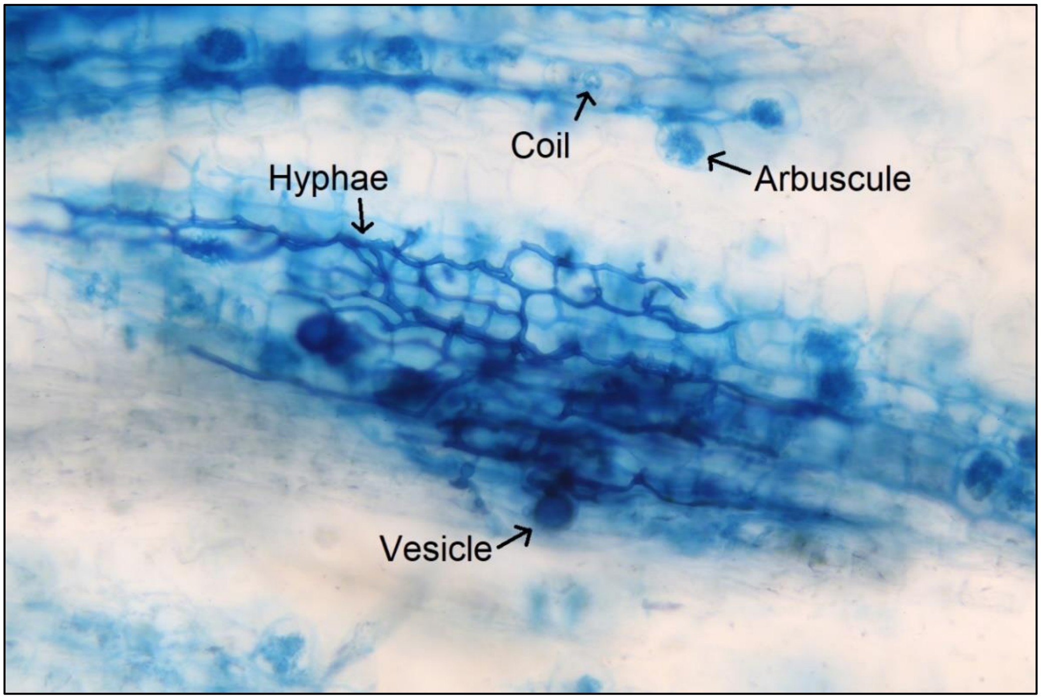


**Figure S1: A micrograph of AM fungal structures quantified in the roots of experimental seedlings.** A photograph taken at 200 x magnification under a compound light microscope shows examples of the four structures quantified in the AM fungal study (hyphae, arbuscules, vesicles, and coils). The structures are stained blue inside the root of an experimental *Virola surinamensis* seedling.

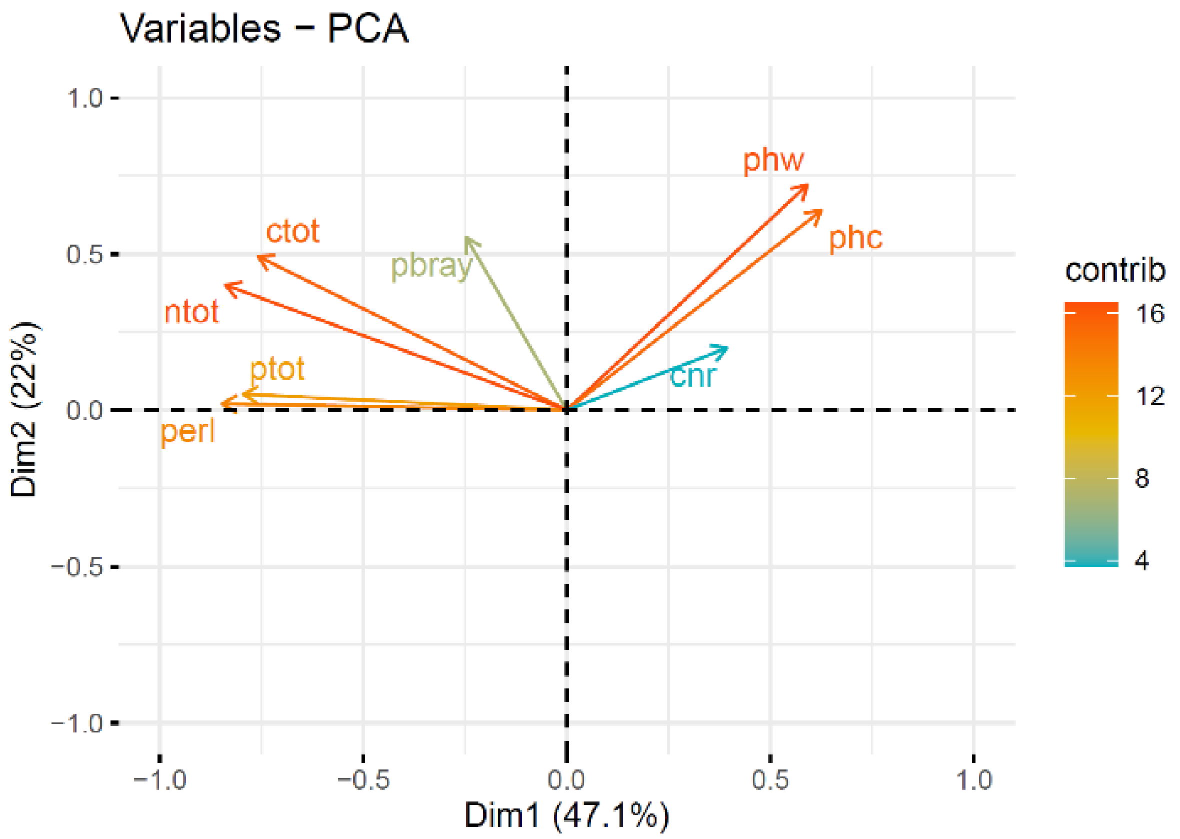


**Figure S2. Loading plot of soil nutrient variable contributions and dimensionality in principal component analysis.** Principal component analysis (PCA) was used to summarize the eight nutrient variables quantified post-experimentally in the 21 soil inocula used in the shadehouse experiment (Table S6). The loadings of each soil nutrient variable on the first two principal components (PC1 & PC2) resulting from PCA are shown. Longer arrow lengths represent higher relative contributions to the component. Soil nutrient abbreviations: cnr = carbon (C) : nitrogen (N) ratio, phc = pH in CaCl2, phw = pH in water, pbray = plant-available phosphorous (P) (Bray 1-P mg/soil kg), ctot = % total C, ntot = % total N, ptot = total P (mg/soil kg), and perl = organic matter (% loss on ignition).

**Table S1: Effect of soil microbial treatment on seedling biomass in the shadehouse experiment.** We examined the effects of soil microbial treatment (female conspecific soil, male conspecific soil, or heterospecific soil) on the final oven-dried biomass of *Virola surinamensis* seedlings that survived until the end of a shadehouse experiment on Barro Colorado Island using a linear mixed-effects model ($n$ = 246 seedlings). Type III analysis of variance results are shown below, including sum of squares (*SS*), mean square (*MS*), numerator degrees of freedom (Num. *d.f.*), denominator degrees of freedom (Den. *d.f.*), F-score ($F$), and p-values ($p$). Effects for which $p < 0.05$ are highlighted in bold.

| Analysis of Variance | *SS* | *MS* | *Num. d.f.* | *Den. d.f.* | *F* | *p* |
|---|---|---|---|---|---|---|
| Soil Microbiota Treatment | 1.55 | 0.77 | 2 | 15.31 | 1.52 | 0.25 |
| Initial Seedling Biomass | 54.71 | 54.71 | 1 | 234.91 | 107.39 | **$< 2.00 \times e^{-16}$** |
| Soil Nutrient Conditions | 0.98 | 0.98 | 1 | 15.72 | 1.93 | 0.18 |
| Adult Tree Size (DBH) | 0.84 | 0.84 | 1 | 13.14 | 1.64 | 0.22 |

**Table S2: Effect of soil microbial treatment on seedling colonization by AM fungi in the shadehouse experiment.** At the end of a shadehouse experiment on BCI, we tested whether the proportion of root intersects colonized by AM fungi varied among *Virola surinamensis* seedlings inoculated with female conspecific soil microbial communities, male conspecific soil microbial communities, and heterospecific soil microbial communities using a generalized linear mixed-effects model ($n$ = 230 seedlings). We test this using all visible AM fungal structures (Table S2A), arbuscules only (Table S2B), and vesicles only (Table S2C). Type II analysis of deviance results from Wald chi-square tests are shown below, including chi-squared (*Chisq*), degrees of freedom (*d.f.*) and p-values ($p$). Effects for which $p < 0.05$ are highlighted in bold.

<u>Table S2A: All visible AM fungal structures</u>

| Analysis of Deviance | *Chisq* | *d.f.* | *p* |
|---|---|---|---|
| Soil Microbiota Treatment | 7.15 | 2 | **0.03** |
| Initial Seedling Biomass | 3.01 | 1 | 0.08 |
| Soil Nutrient Conditions | 0.52 | 1 | 0.47 |
| Adult Tree Size (DBH) | 1.30 | 1 | 0.25 |
| Observer | 39.73 | 1 | **2.92 x e$^{-10}$** |

<u>Table S2B: Arbuscules only</u>

| Analysis of Deviance | *Chisq* | *d.f.* | *p* |
|---|---|---|---|
| Soil Microbiota Treatment | 8.37 | 2 | **0.02** |
| Initial Seedling Biomass | 0.07 | 1 | 0.79 |
| Soil Nutrient Conditions | 0.71 | 1 | 0.40 |
| Adult Tree Size (DBH) | 2.61 | 1 | 0.11 |
| Observer | 58.46 | 1 | **2.08 x e$^{-14}$** |

Table S2 continued on next page.

Table S2C: Vesicles only

| Analysis of Deviance | *Chisq* | *d.f.* | *p* |
|---|---|---|---|
| Soil Microbiota Treatment | 2.50 | 2 | 0.29 |
| Initial Seedling Biomass | 1.83 | 1 | 0.18 |
| Soil Nutrient Conditions | 0.01 | 1 | 0.94 |
| Adult Tree Size (DBH) | 0.04 | 1 | 0.85 |
| Observer | 0.32 | 1 | 0.57 |

**Table S3: Effect of female conspecific, male conspecific, and heterospecific field environments on seedling survival in the field experiment.** Using a generalized linear mixed-effects model, we tested whether the survival of *V. surinamensis* seedlings at the end of a 7-mo field experiment on BCI varied among field environments (female conspecific, male conspecific, or heterospecific; $n$ = 249 seedlings; Table S3A). Due to missing size information for one heterospecific focal tree (H6), we report the effect of adult tree size using a separate model ($n$ = 239 seedlings; Table S3B). Type II analysis of deviance results from Wald chi-square tests are shown below, including chi-squared (*Chisq*), degrees of freedom (*d.f.*) and p-values ($p$). Effects for which $p < 0.05$ are highlighted in bold.

Table S3A: Without effect of adult tree size

| **Analysis of Deviance** | ***Chisq*** | ***d.f.*** | ***p*** |
|---|---|---|---|
| Field Soil Environment | 0.70 | 2 | 0.71 |
| Initial Seedling Biomass | 16.23 | 1 | **5.60 x $e^{-05}$** |
| Seedling Damage by Herbivores | 4.10 | 1 | 0.07 |
| Conspecific Seedling Density | 4.63 | 1 | **0.03** |

Table S3B: With the effect of adult tree size

| **Analysis of Deviance** | ***Chisq*** | ***d.f.*** | ***p*** |
|---|---|---|---|
| Field Soil Environment | 0.75 | 2 | 0.69 |
| Initial Seedling Biomass | 15.68 | 1 | **7.50 x $e^{-05}$** |
| Seedling Damage by Herbivores | 3.99 | 1 | **0.05** |
| Conspecific Seedling Density | 3.64 | 1 | 0.06 |
| Adult Tree Size (DBH) | 0.18 | 1 | 0.67 |

**Table S4: Effect of female conspecific, male conspecific, and heterospecific field environments on seedling biomass in the field experiment.** Using a linear mixed effects model, we tested whether the harvest biomass of *V. surinamensis* seedlings that survived until the end of a 7-mo field experiment on BCI varied among female conspecific, male conspecific, and heterospecific field environments ($n$ = 40 seedlings; Table S4A). Due to missing size information for one heterospecific focal tree (H6), we report the effect of adult tree size using a separate model ($n$ = 37 seedlings; Table S4B). Analysis of variance results are shown below, including sum of squares (*SS*), mean square (*MS*), numerator degrees of freedom (*Num. d.f.*), denominator degrees of freedom (*Den. d.f.*), F-score (*F*), and p-values (*p*).

<u>Table S4A: Without the effect of adult tree size</u>

| **Analysis of Variance** | ***SS*** | ***MS*** | ***Num. d.f.*** | ***Den. d.f.*** | ***F*** | ***p*** |
|---|---|---|---|---|---|---|
| Field Environment | 0.02 | 0.01 | 2 | 16.72 | 0.33 | 0.73 |
| Initial Seedling Biomass | 0.06 | 0.06 | 1 | 35.51 | 1.94 | 0.17 |
| Seedling Damage by Herbivores | 0.79 | 0.79 | 1 | 30.40 | 25.01 | **2.25 x e$^{-05}$** |
| Conspecific Seedling Density | 0.05 | 0.05 | 1 | 32.15 | 1.62 | 0.21 |

<u>Table S4B: With the effect of adult tree size</u>

| **Analysis of Variance** | ***SS*** | ***MS*** | ***Num. d.f.*** | ***Den. d.f.*** | ***F*** | ***p*** |
|---|---|---|---|---|---|---|
| Field Environment | 0.04 | 0.02 | 2 | 15.96 | 0.66 | 0.53 |
| Initial Seedling Biomass | 0.07 | 0.07 | 1 | 30.85 | 2.02 | 0.16 |
| Seedling Damage by Herbivores | 0.65 | 0.65 | 1 | 27.31 | 19.55 | **1.41 x e$^{-04}$** |
| Conspecific Seedling Density | 0.02 | 0.02 | 1 | 26.69 | 0.59 | 0.45 |
| Adult Tree Size (DBH) | 0.04 | 0.04 | 1 | 14.85 | 1.19 | 0.29 |

**Table S5: Effect of female conspecific, male conspecific, and heterospecific field environments on seedling relative growth rates in the field experiment.** Using a linear mixed-effects model, we tested whether the relative growth rates of *V. surinamensis* seedlings during a 7-mo field experiment on BCI varied among female conspecific, male conspecific, and heterospecific field environments ($n$ = 350 RGR observations; Table S5A). Due to missing size information for one heterospecific focal tree (H6), we report the effect of adult tree size using a separate model ($n$ = 331 RGR observations; Table S5B). Type III analysis of variance results are shown below, including sum of squares (*SS*), mean square (*MS*), numerator degrees of freedom (Num. *d.f.*), denominator degrees of freedom (Den. *d.f.*), F-score (*F*), and p-values (*p*).

<u>Table S5A: Without the effect of adult tree size</u>

| **Analysis of Variance** | ***SS*** | ***MS*** | ***Num. d.f.*** | ***Den. d.f.*** | ***F*** | ***p*** |
|---|---|---|---|---|---|---|
| Field Environment | $7.27 \times e^{-05}$ | $3.63 \times e^{-05}$ | 2 | 335.89 | 0.38 | 0.68 |
| Initial Seedling Biomass | $1.41 \times e^{-03}$ | $1.41 \times e^{-03}$ | 1 | 128.79 | 14.91 | **$1.77 \times e^{-04}$** |
| Seedling Damage by Herbivores | $1.90 \times e^{-03}$ | $1.90 \times e^{-03}$ | 1 | 300.13 | 20.14 | **$1.03 \times e^{-05}$** |
| Conspecific Seedling Density | $9.08 \times e^{-05}$ | $9.08 \times e^{-05}$ | 1 | 336.78 | 0.96 | 0.33 |

<u>Table S5B: With the effect of adult tree size</u>

| **Analysis of Variance** | ***SS*** | ***MS*** | ***Num. d.f.*** | ***Den. d.f.*** | ***F*** | ***p*** |
|---|---|---|---|---|---|---|
| Field Environment | $2.44 \times e^{-05}$ | $1.22 \times e^{-05}$ | 2 | 317.27 | 1.32 | 0.27 |
| Initial Seedling Biomass | $8.34 \times e^{-04}$ | $8.34 \times e^{-04}$ | 1 | 115.59 | 9.02 | **$3.27 \times e^{-03}$** |
| Seedling Damage by Herbivores | $1.44 \times e^{-03}$ | $1.44 \times e^{-03}$ | 1 | 278.71 | 15.56 | **$1.01 \times e^{-04}$** |
| Conspecific Seedling Density | $8.28 \times e^{-05}$ | $8.28 \times e^{-05}$ | 1 | 319.99 | 0.90 | 0.35 |
| Adult Tree Size (DBH) | $1.15 \times e^{-06}$ | $1.15 \times e^{-06}$ | 1 | 271.55 | 0.01 | 0.91 |

**Table S6: Summary of principal component analysis of nutrients in the soil inocula at the end of the shadehouse experiment**. We conducted principal component analysis (PCA) on the 21 soil inocula in the shadehouse experiment using data from eight soil nutrient variables. The standard deviation, proportion of variance, and cumulative variance are shown for the first eight principal components resulting from PCA (Table S6A). Loadings of the soil nutrient variables on the principal components resulting from PCA are shown (Table S6B). Soil nutrient variable abbreviations: cnr = carbon (C) : nitrogen (N) ratio, phc = pH in CaCl2, phw = pH in water, pbray = plant-available phosphorous (P) (Bray 1-P mg/soil kg), ctot = % total C, ntot = % total N, ptot = total P (mg/soil kg), and perl = organic matter (% loss on ignition).

Table S6A: Principal components from PCA

| Summary | Comp. 1 | Comp. 2 | Comp. 3 | Comp. 4 | Comp. 5 | Comp. 6 | Comp. 7 | Comp. 8 |
|---|---|---|---|---|---|---|---|---|
| Standard deviation | 1.89 | 1.30 | 1.01 | 0.83 | 0.61 | 0.50 | 0.16 | 0.03 |
| Proportion of variance | 0.47 | 0.22 | 0.13 | 0.09 | 0.05 | 0.03 | < 0.01 | < 0.01 |
| Cumulative variance | 0.47 | 0.69 | 0.82 | 0.91 | 0.96 | 1.00 | 1.00 | 1.00 |

Table S6B: Loadings of the soil nutrient variables on the principal components

| Soil variables | Comp. 1 | Comp. 2 | Comp. 3 | Comp. 4 | Comp. 5 | Comp. 6 | Comp. 7 | Comp. 8 |
|---|---|---|---|---|---|---|---|---|
| phw | 0.31 | 0.56 | 0.24 | - | 0.15 | - | 0.71 | - |
| phc | 0.33 | 0.49 | 0.35 | - | 0.18 | 0.10 | -0.69 | - |
| pbray | -0.13 | 0.43 | -0.48 | 0.68 | -0.21 | 0.22 | - | - |
| ptot | -0.42 | - | - | 0.27 | 0.81 | -0.32 | - | - |
| perl | -0.45 | - | 0.14 | -0.21 | 0.16 | 0.84 | - | - |
| ctot | -0.40 | 0.38 | - | -0.38 | -0.20 | -0.25 | - | -0.67 |
| ntot | -0.44 | 0.31 | 0.13 | -0.17 | -0.29 | -0.26 | - | 0.71 |
| cnr | 0.21 | 0.15 | -0.74 | -0.49 | 0.31 | - | - | 0.22 |

### References Cited Only in the Supplementary Information


- Condit, R., Engelbrecht, B. M. J., Pino, D., Perez, R., & Turner, B. L. (2013). Species distributions in response to individual soil nutrients and seasonal drought across a community of tropical trees. *PNAS* 110(13): 5064-5068.

- Zalamea, P.-C., Turner, B. L., Winter, K., Jones, F. A., Sarmiento, C., & Dalling, J. W. (2016). Seedling growth responses to phosphorus reflect adult distribution patterns of tropical trees. *New Phytologist* 212(2): 400-408.
